# Membrane Mimetic-Dependence of GPCR Energy Landscapes

**DOI:** 10.1101/2023.10.16.562552

**Authors:** Naveen Thakur, Arka Prabha Ray, Edward Lyman, Zhan-Guo Gao, Kenneth A. Jacobson, Matthew T. Eddy

## Abstract

Protein function strongly depends on temperature, which is related to temperature-dependent changes in the equilibria of protein conformational states. We leveraged variable-temperature ^19^F-NMR spectroscopy to interrogate the temperature dependence of the conformational landscape of the human A_2A_ adenosine receptor (A_2A_AR), a class A GPCR. Temperature-induced changes in the conformational equilibria of A_2A_AR in lipid nanodiscs were markedly dependent on the efficacy of bound drugs. While antagonist complexes displayed only modest changes as the temperature rose, both full and partial agonist complexes exhibited substantial increases in the active state population. Importantly, the temperature-dependent response of complexes with both full and partial agonists exhibited a pronounced sensitivity to the specific membrane mimetic employed. In striking contrast to observations within lipid nanodiscs, in detergent micelles the active state population exhibited different behavior for A_2A_AR complexes with both full and partial agonists. This underscores the importance of the protein environment in understanding the thermodynamics of GPCR activation.

**Highlights:**

- Active A_2A_AR population increases with increasing temperature in lipid nanodiscs
- Active A_2A_AR population exhibits different temperature dependence in detergents
- Partial agonist complexes present a unique conformational state in nanodiscs
- Temperature dependence of partial agonist equilibria depends on membrane mimetic

## Introduction

Numerous studies have documented the ability of proteins to adopt multiple conformational states at physiological temperatures and have linked dynamic fluctuations among different conformations with protein biological functions.^1-9^ Some of the first investigations of this connection over forty years ago observed the motion of aromatic residues in globular proteins using NMR spectroscopy, providing valuable insights into enzymatic functions.^10,11^ Subsequently, there has been a continued exploration of the internal dynamics of proteins, spanning various timescales and protein types, using NMR and a growing number of spectroscopic techniques.^12^

A prevailing perspective underscores the significance of dynamics, or structural plasticity, in the functions of G protein-coupled receptors (GPCRs).^13,14^ Structural plasticity enables GPCRs to bind a diverse array of molecules and interact with intracellular partner proteins, leading to the formation of signaling complexes at the plasma membrane. Evidence supporting the existence of multiple, functionally-relevant conformations is robust and draws heavily from the spectroscopic literature, especially NMR spectroscopy.^15-18^ NMR studies have consistently identified the presence of multiple conformations simultaneously coexisting in a functionally relevant equilibrium for an expanding repertoire of GPCRs.^15,18-38^

In previous investigations, we demonstrated how the conformational equilibria of the human A_2A_ adenosine receptor (A_2A_AR), a representative class A GPCR, was influenced not only by the efficacy of bound drugs but also significantly affected by the presence and absence of specific phospholipids.^19,39^ Our ^19^F-NMR data recorded at a single temperature of 280 K distinctly revealed clear disparities in the conformational equilibria of A_2A_AR between detergent and lipid nanodisc preparations.^19^ These studies were conducted within the broader context of a growing interest in exploring the role of lipids in shaping the structure-function relationships of GPCRs.^26,40-47^ Building upon these findings, the current study leverages variable temperature NMR to provide insights into the temperature dependence of A_2A_AR activation and to compare the temperature-dependence of activation across different membrane mimetic systems.

We also expand upon prior research involving A_2A_AR complexes with full agonists by examining the temperature-dependent response of A_2A_AR when bound to partial agonists, which exhibit a spectrum of efficacies spanning between the receptor’s basal state to just under full activity. For complexes with partial agonists, we observe the emergence of a distinct conformational state exclusive to these complexes in lipid nanodiscs. We also discern differences in the response of the conformational equilibria to changes in temperature, which depend on both the efficacy of bound drugs and the characteristics of the membrane mimetic. This data is evaluated within the broader context of understanding the mechanisms underlying GPCR-G protein recognition and is discussed in the context of thermodynamic models of GPCR activation.

## Results

### The population of an active A_2A_AR conformational state increases with increasing temperature for agonist-bound A_2A_AR

For variable temperature NMR experiments, we employed a variant of human A_2A_AR containing a single cysteine replacement at position 289 located at the intracellular surface of TM VII (Figure S1), A_2A_AR[A289C]. This variant was expressed in *Pichia pastoris* using previously described protocols^19,48,49^ and labeled with ^19^F-2,2,2-trifluorethanethiol (TET) at position 289C for NMR experiments following an in-membrane chemical modification approach,^50^ which ensured that position 289 was the only site labeled. The ^19^F-NMR probe introduced at position 289 was shown to be a sensitive reporter of A_2A_AR conformational changes resulting from complex formation with ligands of different efficacies^22^ and resulting from changes in the membrane mimetic environment,^19,39^ thus providing a “fingerprint” of the corresponding A_2A_AR functional states. ^19^F-labeled A_2A_AR was purified and studied in both detergent environments and lipid nanodiscs containing different binary and ternary lipid mixtures following established reconstitution protocols.^19,49^

We recorded 1-dimensional ^19^F-NMR spectra of A_2A_AR[A289C^TET^] in complex with the full agonist 5’-N-ethylcarboxamidoadenosine (NECA) reconstituted in lipid nanodiscs containing 1-palmitoyl-2-oleoyl-glycero-3-phosphocholine (POPC) and 1-palmitoyl-2-oleoyl-sn-glycero-3-phospho-L-serine (POPS), at a 70:30 molar ratio, at three temperatures: 280 K, 298 K, and 310 K. Spectra of A_2A_AR in complex with NECA measured at 280 K were consistent with previously published data at the same temperature.^19,39^ At 280 K, we observed four distinct states labeled P1, P2, P4, and P5, with P4 showing the largest signal intensity (Figure 1). In previously published studies, states P2 and P4 had been observed only for agonist-bound A_2A_AR.^19,22^ State P4 was assigned previously to a fully active A_2A_AR conformation by carefully conducting experiments in which we observed a population of P4 only upon complex formation of A_2A_AR[A289C^TET^] with mini-G_S_^19^, an engineered G protein.^51^ Deconvolutions of agonist-bound A_2A_AR in previous studies did not include P5, because P5 could only be confidently identified in spectra of A_2A_AR bound to partial agonists in nanodiscs (see below), which allowed us to refit data for agonist-bound A_2A_AR using a refined 5-state model. The presence of states P1, P2, and P4 for agonist-bound A_2A_AR was detected over the entire range of temperatures, but there was a clear temperature-dependent variation in the relative signal intensities. As the integrated signal intensities are proportionate to the relative populations of each conformational state, we thus observed a shift in the conformational equilibria as a function of increasing temperature. The relative population of P4 increased successively from 44% of the total integrated signal intensity at 280 K, to 47% at 298 K, and further to 68% at 310 K (Figure 1 and Table S1). This ∼54% increase in the relative population of P4 over the range of temperatures was accompanied by decreases in the relative populations of both P2 and P1 with increasing temperature (Figure 1A and 1C). With increasing temperature, we observed line narrowing likely due to faster molecular tumbling at higher temperatures, and small upfield shifts of the chemical shifts for all states, consistent with similar observations of temperature dependent upfield shifts reported in earlier ^19^F-NMR studies of β_2_AR.^52^

**Figure 1.**
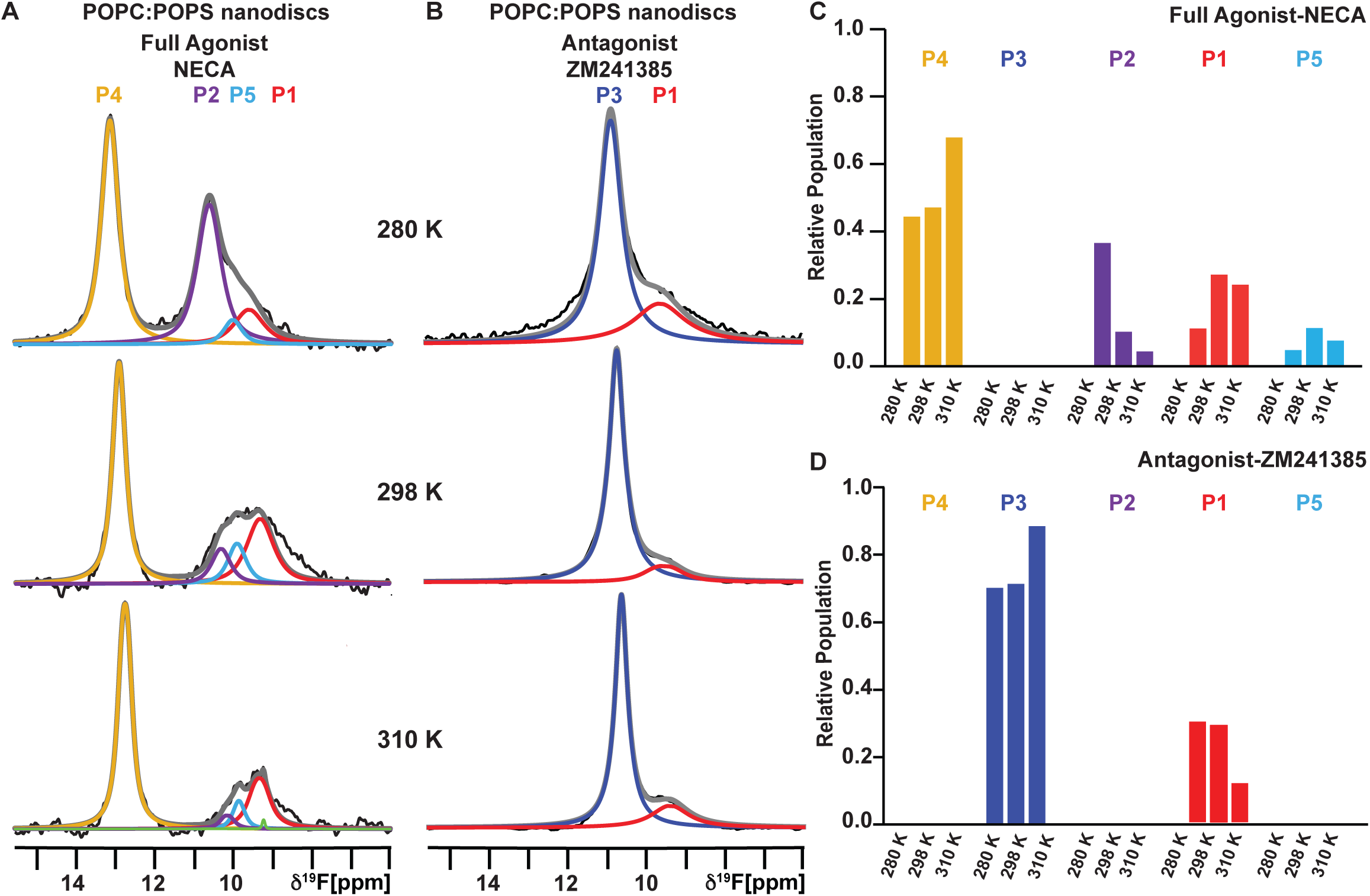
^19^F-NMR observed conformational ensemble of A_2A_AR-ligand complexes in lipid nanodiscs compared over variable temperatures. (A and B) The 1-dimensional ^19^F-NMR spectra of A_2A_AR[A289C^TET^] reconstituted into lipid nanodiscs containing POPC:POPS (70:30 molar ratio) recorded at 280 K, 298 K and 310 K for complexes with (A) the full agonist NECA and (B) the antagonist ZM241385. The NMR spectra are interpreted by Lorentzian deconvolutions with the minimal number of components that provided a good fit, labeled P1 to P4, superimposed on the frequency-domain data (black lines). Summation of the individual fits is shown by the superimposed grey lines. The green fitted line at 310 K is from an NMR signal from free ^19^F-TET. (C and D) Relative populations of each state over the range of studied temperatures determined from the fitted data shown in A and B represented in a bar chart format for A_2A_AR in complex with the (C) agonist NECA and (D) antagonist ZM241385. See also Figures S1 – S4 and Table S1.

We extended the variable temperature study of agonist-bound A_2A_AR in nanodiscs containing POPC and POPS to agonist-bound A_2A_AR in nanodiscs containing binary mixtures of POPC and either PtdIns(4,5)P_2_ (PI(4,5)P_2_), or 1-palmitoyl-2-oleoyl-sn-glycero-3-phosphate (POPG), and agonist-bound A_2A_AR in nanodiscs containing ternary mixtures of POPC, POPS or POPG, and cholesterol. Similar to the temperature-dependent behavior of A_2A_AR[A289C^TET^] in nanodiscs containing POPC and POPS, we observed a significant increase in the relative population of state P4 with increasing temperature with these additional binary and ternary lipid mixtures (Figure S2; Table S1). For agonist-bound A_2A_AR in nanodiscs containing POPC and PI(4,5)P_2_ (95:5, molar ratio), the relative population of the active state P4 increased from 51% at 280 K to 74% at 298 K (Figure S2; Table S1), and for agonist-bound A_2A_AR in nanodiscs containing POPC and POPG (70:30, molar ratio), the relative population of P4 increased from 43% at 280K to 70% at 298K (Figure S2; Table S1). Additionally, agonist-bound A_2A_AR in nanodiscs containing cholesterol exhibited an increase in the relative population of P4 (Figure S2; Table S1). Thus, for agonist-bound A_2A_AR, in all nanodiscs containing binary mixtures of zwitterionic and anionic phospholipids, and in ternary mixtures of phospholipids and cholesterol, we consistently observed an increase in the active state population and subsequent decrease in the populations of other states.

### The conformational ensemble of antagonist-bound A_2A_AR is insensitive to temperature

To assess the temperature response of the antagonist-bound A_2A_AR conformational ensemble, we recorded 1-dimensional ^19^F-NMR spectra of A_2A_AR[A289C^TET^] in nanodiscs containing POPC and POPS (70:30 molar ratio) in complex with the antagonist ZM241385 at 280 K, 298 K and 310 K (Figure 1). For antagonist-bound A_2A_AR at 280 K, we observed two populations, P1 and P3, with P3 being the significantly more intense signal, consistent with earlier results in lipid nanodiscs^19^ and in DDM/CHS micelles.^22^ In contrast to agonist-bound A_2A_AR, the relative populations of states P3 and P1 were at most marginally affected by changing temperature (Figure 1 and Table S1). Line narrowing and small upfield shifts were observed with increasing temperatures, consistent with increasing frequency of rotational motion, but the relative populations of P3 and P1 remained highly similar at all temperatures. Notably, we did not observe any detectable population for the active state P4, even at the highest recorded temperature.

### A_2A_AR complexes with partial agonists manifest a unique conformational state

To investigate the temperature dependence of A_2A_AR activation across the complete range of ligand efficacies, we recorded 1-dimensional ^19^F-NMR data of A_2A_AR in lipid nanodiscs containing POPC and POPS (70:30 molar ratio) for each of two complexes with the partial agonists nonnucleoside LUF5834^53,54^ and nucleoside Regadenoson, an FDA-approved drug used to evaluate heart conditions.^55,56^ The 1-dimensional spectrum of A_2A_AR[A289C^TET^] in complex with Regadenoson recorded at 280 K showed 4 discernible components, including state P4, previously observed in spectra of full agonist complexes, state P3 observed in spectra of antagonist complexes, and state P1, which appeared in spectra of both antagonist and agonist complexes. Additionally, we observed a new component, labeled as P5, with a chemical shift of 10.05 ppm (Figure 2A). Notably, this signal had not been detected previously in spectra of apo, agonist-bound, or antagonist-bound A_2A_AR but was observed in spectra of the complex with Regadenoson at all recorded temperatures (Figure 2A). Deconvolution of the ^19^F-NMR spectra of Regadenoson-bound A_2A_AR using a 4-state model that initially excluded P5 resulted in significant non-zero remaining signal intensity at the chemical shift of P5 (Figure S3), supporting the use of a 5-state model for deconvolution of spectra of A_2A_AR in complex with Regadenoson. The chemical shift of component P5 was distinct from reported chemical shifts for free TET, which may be present at trace amounts in ^19^F-labeled GPCR samples.^57^ The linewidth (FWHM) for component P5 for the complex with Regadenoson was 370 Hz at 280 K, comparable to the line widths for the other observed signals, further supporting the conclusion that the signal P5 originates from the receptor rather than free TET.

**Figure 2.**
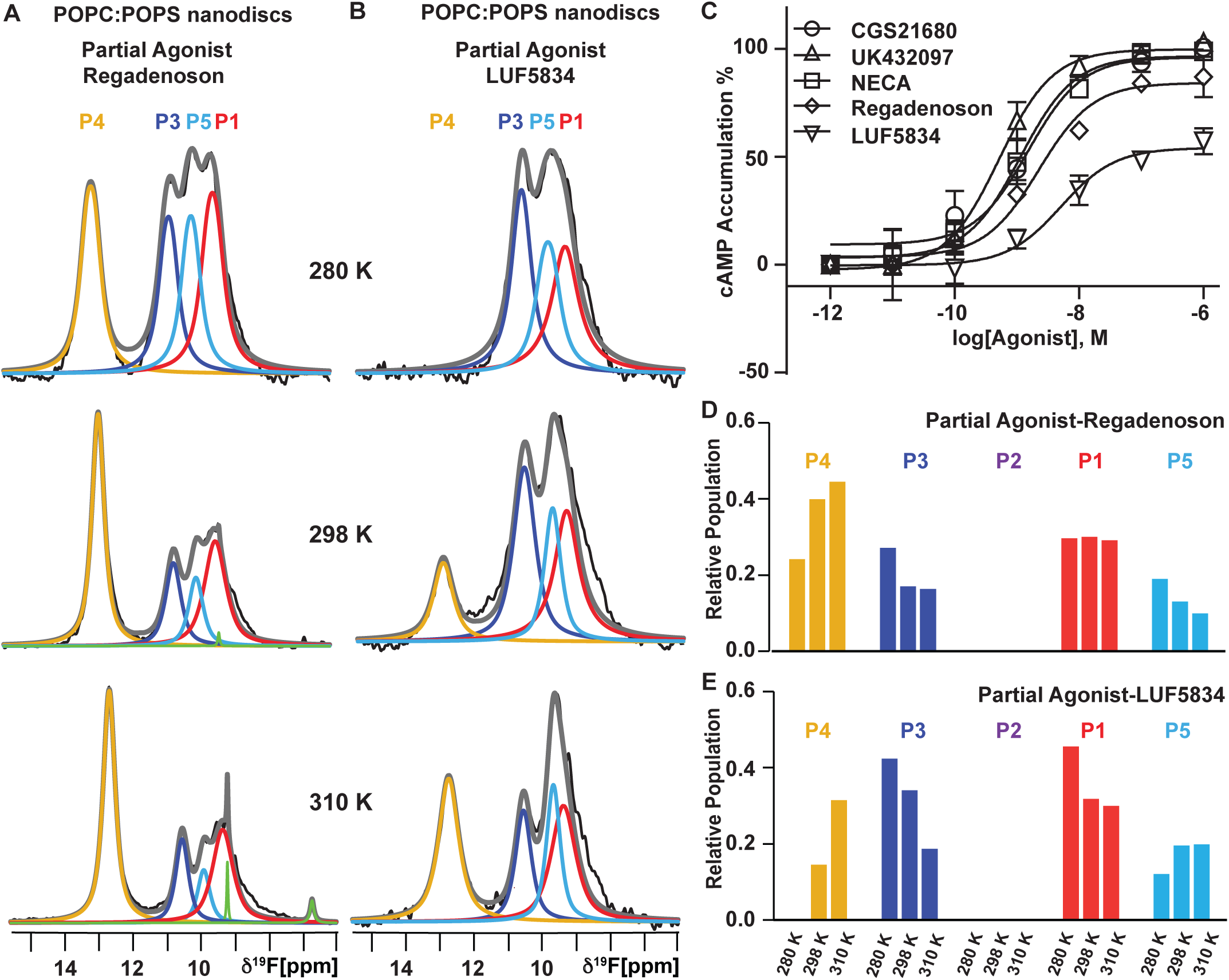
Variable temperature ^19^F-NMR observed ensembles for complexes with partial agonists of differing efficacy. (A and B) ^19^F-NMR spectra of A_2A_AR[A289C^TET^] in lipid nanodiscs containing POPC:POPS (70:30 molar ratio) recorded at three temperatures for complexes with the partial agonists (A) Regadenoson and (B) LUF5834. The fit for the unique population observed for the partial agonist complexes is shown by the cyan line. Other figure presentation details the same as in Figure 1. (C) cAMP accumulation experiments upon stimulation of A_2A_AR with full and partial agonists. Data are shown as the means ± standard error of mean (s.e.m.) for three independent experiments. The EC_50_ values are 1.52 nM, 1.14 nM, 0.52 nM, 5.14 nM, and 2.26 nM for CGS21680, NECA, UK432097, LUF5834, and Regadenoson, respectively. The E_max_ values were 101%, 100%, 103%, 57%, and 87% for CGS21680, NECA, UK432097, LUF5834, and Regadenoson, respectively. (D and E) Relative populations of each state over the range of studied temperatures determined from the fitted data shown in A and B in a bar chart format for A_2A_AR in complex with the partial agonist (D) Regadenoson and (E) LUF5834. See also Figures S1 – S3 and Table S1.

The component P5 was also observed in 1-dimensional ^19^F-NMR spectra of A_2A_AR[A289C^TET^] in complex with a second partial agonist LUF5834 (Figure 2B) at the same chemical shift as observed spectra of A_2A_AR bound to Regadenoson. While the signal corresponding to P5 was less resolved compared to the complex with Regadenoson, deconvolution of the spectra confirmed that a good overall fit could only be obtained by using the same 5-state model utilized for interpreting the spectra for the complex with Regadenoson (Figure S3). While we did incorporate P5 into the deconvolution of NMR data for A_2A_AR[A289C^TET^] in complex with the full agonist NECA (Figure 1), it did not significantly change the overall fit (Figure S3), nor was P5 discernible in spectra of A_2A_AR[A289C^TET^] in complex with the antagonist ZM241385. Consequently, the presence of a significant population for the P5 conformational state appears to be unique to complexes involving partial agonists in lipid nanodiscs. This finding underscores that earlier ^19^F-NMR studies of A_2A_AR complexes with full agonists and antagonists in lipid nanodiscs ^19,39^ would not have been affected by the observation of the P5 signal, as it appears to be significantly populated only for partial agonist complexes.

### The population of active A_2A_AR increases with increasing temperature for complexes with partial agonists

At 280 K, the ^19^F-NMR spectra recorded with LUF5834 and Regadenoson displayed notable differences in the relative peak intensities, with the most striking difference observed in the signal P4. For the complex with Regadenoson, the integrated signal intensity for P4 accounted for approximately 24% of the total integrated signal intensity. In contrast, for the complex with LUF5834, the signal corresponding to peak 4 was not detected at all (Figure 2A and Figure 2B; Table S1). For the complex with Regadenoson, the relative integrated signal intensity of P4 exhibited a temperature-dependent increase, rising from ∼24% at 280 K to 40% at 298 K, and reaching 45% at 310 K. This increase was accompanied by a simultaneous decrease in the relative intensities of states P3, P5 and P1 with increasing temperature (Figure 2A and 2D; Table S1). Comparatively, when contrasted with the complex involving the full agonist NECA, the signal intensity of P4 for the Regadenoson complex was approximately 51% lower at 310 K (Table S1).

In line with our observations of the temperature-dependent response of the complex with Regadenoson, we also observed a temperature-dependent change in the relative populations of each observed state for the complex with LUF5834. Upon increasing the temperature from 280 K to 298 K, we observed the emergence of signal P4, present at ∼15% of the total integrated signal intensity (Figure 2B and 2E; Table S1). Further increasing the temperature produced a notable increase in the integrated signal intensity for P4, reaching ∼31% of the total integrated signal at 310 K. The increase in P4 coincided with a decrease in the population P3, previously assigned as an inactive state in spectra of antagonist complexes. When comparing the spectra of complexes with Regadenoson and LUF5834 at 310 K, we observed a distinct difference in the relative population of P4, with a lower population of P4 for the complex with LUF5834, 31% for the complex with LUF5834 versus 45% for the complex with Regadenoson. Concurrently, the signal intensity for P5, the state unique to partial agonist complexes, was larger for LUF5834 than Regadenoson, 20% versus 10%, respectively.

The relative differences in the populations for the active state P4 and for the state P5 unique to partial agonist complexes align with findings from cellular signaling assays measuring cAMP accumulation upon A_2A_AR stimulation (Figure 2C). In the cAMP accumulation assays, we observed an efficacy for LUF5834 of ∼50% maximal efficacy compared to full agonists (NECA, CGS21680 and UK432097). For Regadenoson, we observed an efficacy of ∼90% of that for full agonists. The relative differences for the two partial agonists appeared to align with the relative differences in the populations of P4 and P5 between the two partial agonist complexes.

### Temperature-dependent changes in the A_2A_AR conformational ensemble depend on the membrane environment

In an earlier NMR study of A_2A_AR, we demonstrated that at a temperature of 280 K, the relative populations of conformational states varied significantly depending on the specific membrane mimetic system employed in ^19^F-NMR experiments. Specifically, we observed a significantly higher intensity for the active state P4 for agonist complexes in nanodiscs containing binary mixtures of POPC and POPS as compared to agonist complexes in mixed micelles composed of the detergent n-dodecyl-β-D-maltoside (DDM) and cholesteryl hemisuccinate (CHS).^19^ These earlier observations motivated us to investigate whether temperature-dependent changes in the relative populations of A_2A_AR conformational states also varied among different membrane mimetic systems.

We recorded ^19^F-NMR spectra of A_2A_AR[A289C^TET^] in complex with the full agonist NECA in DDM/CHS micelles at three temperatures: 280 K, 298 K, and 310 K (Figure 3A and S4). At 280 K, the conformational ensemble of A_2A_AR[A289C^TET^] showed a more prominent signal P1 and two relatively smaller components P2 and P4. This was consistent with previous publications and distinct from the relative intensities of each conformational state observed in ^19^F-NMR spectra of A_2A_AR in nanodiscs containing POPC and POPS (Figure 1). In contrast to the temperature-dependent behavior of A_2A_AR in nanodiscs, we observed a relatively flat population of P4 as the temperature was increased, from 10% at 280 K to 13% at 310 K (Figure 3A and Figure 3E; Table S1). This stands in striking contrast to the significant increase in the P4 population observed for A_2A_AR in nanodiscs containing POPC and POPS (Figure 1). For agonist-bound A_2A_AR in DDM/CHS, we also observed an increase in the relative population of state P1, from 50% at 280 K to 69% at 310 K (Figure 3A and Figure 3E; Table S1), also in contrast to observations of A_2A_AR in lipid nanodiscs (Figure 1).

**Figure 3.**
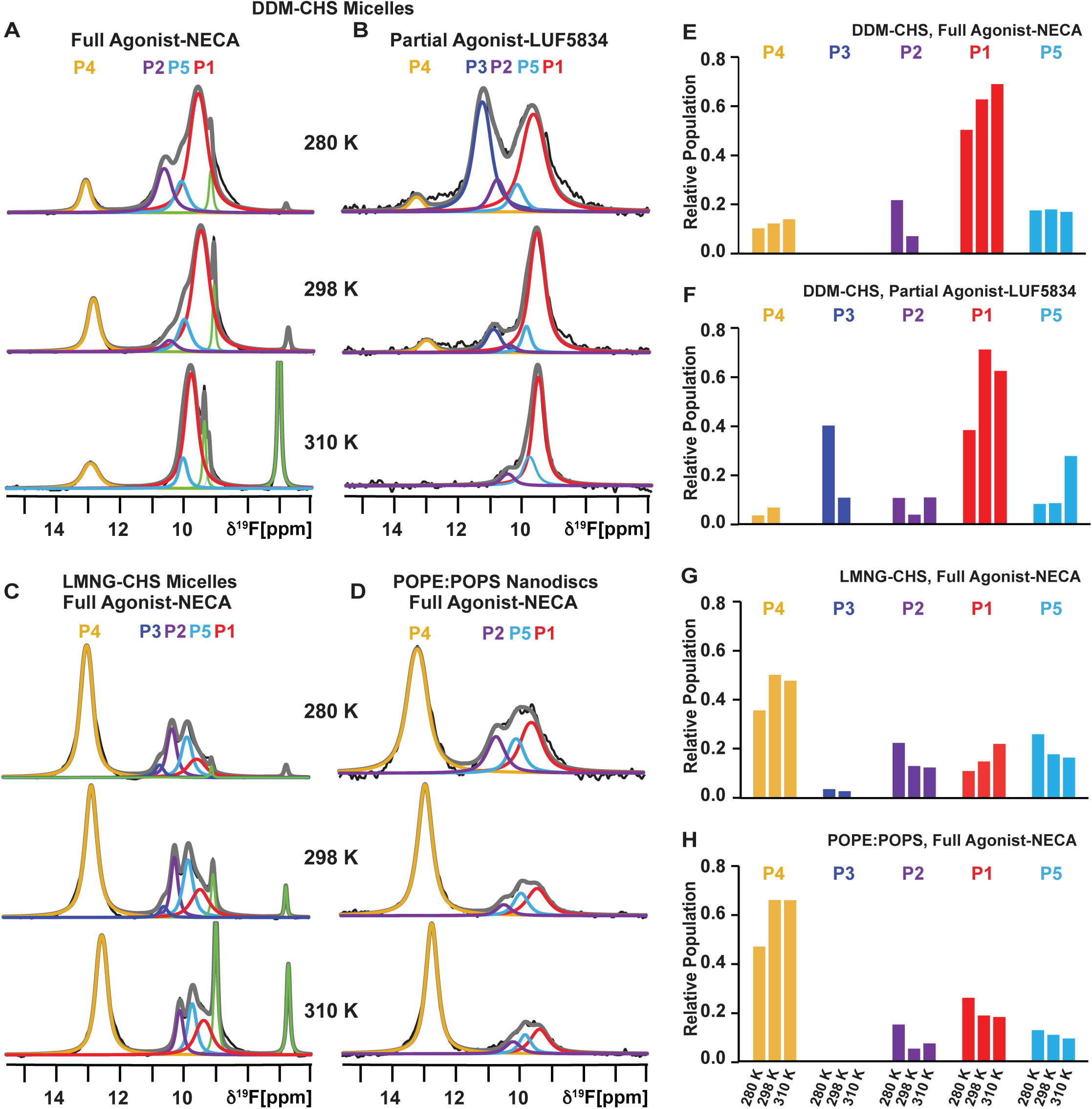
Comparison of the variation in temperature observed for the conformational equilibria of A_2A_AR[A289C^TET^] in three different membrane mimetics. (A and B) Variable temperature ^19^F-NMR spectra of A_2A_AR[A289C^TET^] reconstituted in DDM-CHS micelles in complex with the (A) agonist NECA and (B) partial agonist LUF5834. Other figure presentation details are same as in Figures 1 and 2. (C) Variable temperature ^19^F-NMR spectra of A_2A_AR[A289C^TET^] in complex with the agonist NECA reconstituted in LMNG-CHS micelles. Other figure presentation details are same as in Figures 1 and 2. (D) Variable temperature ^19^F-NMR spectra of A_2A_AR[A289C^TET^] in complex with the agonist NECA reconstituted in POPE:POPS (70:30 molar ratio). Other figure presentation details are same as in Figures 1 and 2. (E – H) Relative populations of each state over the range of studied temperatures determined from the fitted data shown in A – D represented in a bar chart format for A_2A_AR reconstituted in DDM-CHS micelles, in complex with the (E) agonist NECA and (F) partial agonist LUF5834, and A_2A_AR in complex with the agonist NECA reconstituted in (G) LMNG-CHS micelles and (H) lipid nanodiscs containing POPE:POPS (70:30 molar ratio). See also Figures S1 – S4 and Table S1.

We further explored the different responses to temperature between detergent and lipid environments by recording ^19^F-NMR of A_2A_AR[A289C^TET^] in complex with the partial agonist LUF5834 in DDM/CHS micelles. At 280 K, we observed a conformational ensemble for the complex with LUF5834 in DDM/CHS largely dominated by populations P3 and P1, with almost no observable population for P5 in contrast to observations of the complex with LUF5834 in lipid nanodiscs. Another notable difference was the presence of a small but detectable population for the active state P4 in DDM/CHS micelles (Figure 3B). Upon increasing the temperature, we observed that the small population of P4 decreased to a level that was not detectable at 310 K, with a concurrent decrease in the population P3 and increase in the population P1 (Figure 3B and 3F; Table S1).

We extended our ^19^F-NMR comparison to a complex of A_2A_AR[A289C^TET^] with the full agonist NECA in mixed micelles composed of lauryl maltose neopentyl glycol (LMNG) and CHS. Interestingly, for ^19^F-NMR spectra of A_2A_AR agonist complex in LMNG/CHS measured at 280 K, we observed a more complicated signal envelop that contained all 5 components, with component P4 having the largest signal intensity (Figure 3C). Notably, we also observed a significant population for P5, which we had detected for partial agonist complexes in lipid nanodiscs. Upon increasing the temperature, we observed only marginal changes in the relative populations of each conformational state from 280 K to 310 K (Figure 3C and 3G; Table S1).

We also explored the effect of temperature across different lipid environments to explore the response of A_2A_AR to the temperature variance. We recorded 1-dimensional ^19^F-NMR spectra for agonist-bound A_2A_AR in nanodiscs containing the zwitterionic lipid POPE and anionic lipid POPS (Figure 3D). We observed a similar behavior as for agonist-bound A_2A_AR reconstituted in nanodiscs containing POPC and POPS, with P4 being the major population in the ensemble. We also observed a similar temperature-dependent response for agonist-bound A_2A_AR in nanodiscs containing POPE and POPS, showing an increase in relative population of active state P4 with increasing temperature from 280 K to 310 K (Figure 3D and Figure 3H; Table S1).

## Discussion

The NMR spectra shown in Figure 1 reveal clear differences in the temperature response between A_2A_AR complexes with antagonists and agonists. In lipid nanodiscs containing the A_2A_AR complex with the agonist NECA, we observed a distinct trend: a rise in the population of state P4 as the temperature increased, accompanied by a simultaneous decrease in the population of all other states (Figure 1). State P4 had been previously assigned to an A_2A_AR conformation observed for the active receptor in a ternary complex with G protein.^19^ Consequently, we conclude increasing temperature favors a pronounced shift in the conformational ensemble toward increasing the population of this active state. Earlier pharmacological studies of A_2A_AR in membranes noted an increase in the affinity of agonists for A_2A_AR with increasing temperature.^58^ In our current study, all samples were prepared with a saturating concentration of ligand, ensuring that the increase in the active state P4 with increasing temperature was not attributable to differences in affinities at varying temperatures. Nevertheless, our data do suggest a correlation between the increased population of the active state P4 and the observed increase in agonist affinity with increasing temperature. It is also interesting to speculate on correlations between the temperature-dependence of NMR-observed activation and the thermodynamics of ligand binding. It has been proposed that agonist binding is entropy-driven while antagonist-binding is mostly enthalpy driven.^58,59^ This could potentially be related to ligand binding modes: A_2A_AR agonists require hydrophilic interactions deep in the orthosteric binding pocket, while antagonist interactions can be completely hydrophobic. Given the similar ligand binding location and shared chemical scaffolds for agonists and antagonists among adenosine receptor subtypes, A_2A_AR can serve as a guide to predict thermodynamic behavior of other adenosine receptors.

The response observed in agonist-bound A_2A_AR in lipid nanodiscs differs notably from the findings in earlier variable temperature ^19^F-NMR studies of agonist-bound β_2_AR. In a previous ^19^F-NMR study of β_2_AR study, two distinct populations were observed when β_2_AR was labeled with TET at a cysteine on the intracellular surface on either helix VI or helix VII.^60^ These populations were attributed to an inactive state and an active-like state, which were consistently observed in both agonist and antagonist complexes.^60^ Notably, for agonist-bound β_2_AR, the relative population of the active-like conformation decreased as the temperature increased, while the relative population of the inactive state increased.^52^ This shift was significant to the extent that at the highest temperature investigated (310 K), the population of active-like β_2_AR was markedly lower than that of the inactive conformation.^52^

One critical distinction between the β_2_AR study and our current examination of A_2A_AR lies in the employed membrane mimetics. In the β_2_AR study, the receptor was in DDM/CHS micelles, whereas our observations of the increasing active population of A_2A_AR were made in lipid nanodiscs containing binary or ternary mixtures of lipids. For two commonly employed detergent systems, mixed micelles of DDM/CHS and LMNG/CHS, we observed a significant difference in the response to temperature compared with lipid nanodiscs. For agonist-bound A_2A_AR in DDM/CHS micelles, we noted a modest decline in the relative population of active A_2A_AR as the temperature increased (Figure 3). While the magnitude of this decline was not entirely consistent with the more substantial decrease in the active-like state observed for β_2_AR in DDM/CHS from the study by Horst et al.^52^, it was closer in alignment with the β_2_AR results than with the temperature-dependent behavior of A_2A_AR in lipid nanodiscs. When examining agonist-bound A_2A_AR in LMNG/CHS micelles, we observed a relatively higher population of the active state P4 at 280 K compared to DDM/CHS, but this population showed only marginal changes as the temperature increased (Figure 3). From a comparison of the presented data, it is clear that the choice of membrane mimetic significantly influences temperature-dependent changes in the receptor conformational populations. Moreover, this choice at least partially accounts for the significant differences observed between the temperature-dependent response of A_2A_AR in lipid nanodiscs and the earlier study of β_2_AR in detergent micelles. Given that physiological temperatures are closer to 310 K than 280 K, these differences in the temperature response between different membrane mimetics further support the notion that nanodiscs offer a more physiologically relevant environment, provided that the lipid composition is carefully chosen.^19,39^

A substantial body of literature provides compelling evidence that variations in the physical properties of detergent micelles and lipid bilayer environments impact the conformational equilibria of membrane proteins.^61^ This effect is exemplified in the extensive body of work on rhodopsin, where spectroscopic observations have revealed significant disparities in rhodopsin conformational equilibria between detergents and lipid bilayers.^62-65^ These findings prompted development of a flexible surface model that describes the interplay between membrane curvature and hydrophobic forces and membrane protein structure-function relationships.^66,67^ While it is established that an increase in temperature induces negative spontaneous curvature for lipids in vesicles, thereby influencing the conformational equilibria of membrane proteins, the impact on the spatial properties in nanodiscs is less understood but merits exploration in future studies. Additionally, an entropy-favoring activation via hydration has been documented for rhodopsin in lipid bilayers.^68^ It is intriguing to speculate whether differences in water penetration between detergent micelle and lipid bilayer preparations may contribute to variations in GPCR activation.

In our ^19^F-NMR data recorded at 280 K in lipid nanodiscs, we detected a prominent component labeled as ‘P5’ for A_2A_AR when bound to partial agonists (Figure 2). This component had not been previously reported in ^19^F spectra of agonist-bound A_2A_AR when labeled with TET at position A289C, because it is present as only a very minor population and the addition of P5 to fitting the data did not significant change the residual difference between the raw data and overall fit (Figure S3). This observation aligns with findings from 2D NMR correlation studies of A_2A_AR measured at 310K, which reported a unique conformational state for the conserved P-I-F motif and intracellular surface that correlated with the efficacy of partial agonists.^20^ We attributed the existence of P5 to a conformational state that manifested for A_2A_AR within lipid nanodiscs upon complex formation with partial agonists. Furthermore, the relative populations of P5 and P4 for the two complexes with partial agonists LUF5834 and Regadenoson are consistent with results from cellular signaling assays (Figure 2).

The observation of a unique conformational state associated with partial agonism had been reported in earlier ^19^F-NMR studies of A_2A_AR labeled at position V229C in LMNG/CHS micelles, assigned as state ‘S3’.^24^ A notable distinction with our present study is that, for A_2A_AR in lipid nanodiscs, we observed a significant population for the partial agonist conformation only when partial agonists were bound. In our present study, for agonist-bound A_2A_AR in LMNG/CHS micelles, we observed the simultaneous population of all five conformational states, including P5, which appears more in line with previous findings. This observation appears more in line with data from Reference 22 but deviates from our findings for agonist-bound A_2A_AR in lipid nanodiscs containing POPC and POPS, where only three conformational states are observed (Figure 1). This suggests that proposed mechanisms of molecular activation, i.e. conformational selection or induced fit, may also depend on the employed membrane mimetic.

For partial agonist-bound A_2A_AR, we also observed a notable difference in the temperature response between A_2A_AR in DDM/CHS micelles and in lipid nanodiscs (Figure 3). In lipid nanodiscs we observed an increase in the relative population of the active state P4 with rising temperature, however, in detergent micelles, we observed a decrease in P4 to the point where it was no longer observable at 310 K. These findings remain consistent with our observations of agonist-bound A_2A_AR in lipid nanodiscs and further underscore the importance of considering the membrane mimetic in temperature-dependent studies of receptor conformational equilibria.

## Acknowledgements

This work is supported by the National Institutes of Health, NIGMS MIRA grant R35GM138291 (M.T.E., N.T., A.P.R.) and grant number R01GM120351 (E.L.). A portion of this work was supported by the McKnight Brain Institute at the National High Magnetic Field Laboratory’s AMRIS Facility, which is funded by National Science Foundation Cooperative Agreement No. DMR-1644779 and the State of Florida and also supported in part by an NIH award, S10 OD028753, for magnetic resonance instrumentation. The authors also acknowledge support from the NIH/NIDDK Intramural Research Program (ZIA DK031117).

## Author Contributions

M.T.E. designed the study. N.T. and A.P.R. performed protein production, purification, and nanodisc sample preparation. A.P.R. and N.T. recorded NMR data and fluorescence data with input from M.T.E. A.P.R. and N.T analyzed NMR and fluorescence data with input from M.T.E. E. L. provided input on thermodynamic analysis. Z.G.G and K.A.J performed and analyzed cell-based assays. M.T.E., A.P.R. and N.T. wrote the manuscript with input from all authors.

## STAR Methods

### RESOURCE AVAILABILITY

#### Lead Contact

Lead Contact: Dr. Matthew Eddy. Further information and requests for resources or reagents should be directed to and will be fulfilled by the Lead Contact.

#### Materials Availability

All unique reagents generated in this study are available from the Lead Contact by request and with a completed Materials Transfer Agreement.

#### Data and Code Availability

All data reported in this paper and any additional information required to re-analyze the data will be shared by the lead contact upon request. This paper does not report any original code. Any additional information required to reanalyze the data reported in this paper is available from the lead contact upon request.

### EXPERIMENTAL MODEL AND SUBJECT DETAILS

#### Microbes

XL-10 *E. coli* cells were cultured in LB media, BL21(DE3) cells were cultured in TB media, and the BG12 strain of *P. pastoris* was cultured in BMGY and BMMY media.

#### Cell lines

All cell lines used in this study were authenticated by the suppliers and were chosen so as to remain consistent with previously published studies.

### METHOD DETAILS

#### Construct Design

The gene encoding human A_2A_AR (1-316) was cloned into a pPIC9K vector (Invitrogen) at the BamHI and NotI restriction sites. The gene contained a single amino acid replacement (N154Q) to remove the only glycosylation site in the receptor, an N-terminal FLAG tag, and a 10X C-terminal His tag. The construct design was consistent with previous studies.^19,39,49^ No new constructs were generated for this study.

#### A_2A_AR production

Plasmids containing A_2A_AR were transformed into competent cells of the BG12 strain of *Pichia pastoris* via electroporation. High-expressing clones were selected via small-scale protein expression and screening methods where protein expression was detected by an anti-FLAG western blot assay as previously reported. Glycerol stocks of the high expressing clones were stored at -80 °C for future use.

A_2A_AR was expressed in *P. pastoris* following previously published protocols. 4 mL cultures in buffered minimal glycerol (BMGY) media were inoculated from glycerol stocks of a high expressing clone and grown at 30 °C for 48 h. These cultures were consequently used to inoculate 50 mL of BMGY medium and grown at 30 °C for 60 h. Each 50 mL culture was then used to inoculate 500 mL of BMGY medium and then grown for 48 h at 30 °C. The cells were then harvested by centrifugation and then resuspended in 500 mL of buffered minimal methanol (BMMY) media without methanol. The cells were starved for 6 h at 28 °C to consume any remaining glycerol before induction of protein expression by addition of methanol to a final concentration of 0.5% (w/v). Two aliquots of methanol were further added at 12 h interval after induction for a total expression time of 36 h. The cells were then harvested by ultracentrifugation and the cell pellets were frozen in liquid nitrogen and stored at -80 °C for future use.

#### A_2A_AR purification and ^19^F labelling

Purification and ^19^F-labeling via in-membrane chemical modification approach was carried out according to previously published protocols.^49,50^ Cell pellets were resuspended in lysis buffer (50 mM sodium phosphate pH 7.0, 100 mM NaCl, 5% glycerol (w/v), and in-house prepared protease inhibitor solution) and lysed using a cell disruptor (Pressure Biosciences) at 40k PSI. Cell membranes containing A_2A_AR were isolated by ultracentrifugation at 200,000 x g for 30 min and homogenized in buffer (10 mM HEPES pH 7.0, 10 mM KCl, 20 mM MgCl_2_, 1 M NaCl, 4 mM theophylline). The homogenized membranes were incubated with 1 mM of 4,4’-dithiodipyridine (aldrithiol-4) and protease inhibitor cocktail solution for 1 h at 4 °C. The resuspended membranes were pelleted by ultracentrifugation at 200,000 x g for 30 min to remove the excess aldrithiol-4. The pelleted membranes were again resuspended in the same buffer and incubated with 1 mM of TET for 1 h at 4 °C. The membranes were pelleted again using ultracentrifugation, resuspended in the same buffer, and incubated with 1 mM theophylline for 30 min at 4 °C. The protein was extracted by mixing the resuspended membranes 1:1 (v/v) of solubilization buffer (50 mM HEPES pH 7.0, 500 mM NaCl, 0.5% (w/v) n-Dodecyl-β-D-Maltopyranoside (DDM), and 0.05% cholesteryl hemisuccinate (CHS)) for 6 h at 4 °C. The insolubilized material was separated by ultracentrifugation at 200,000 × g for 30 min, and the supernatant was incubated overnight with Co^2+^-charged affinity resin (Talon) and 30 mM imidazole at 4 °C.

After overnight incubation, the resin was washed with 20 CV of wash buffer 1 (50 mM HEPES pH 7.0, 10 mM MgCl_2_, 30 mM imidazole, 500 mM NaCl, 8 mM ATP, 0.05% DDM, and 0.005% CHS), and twice with 20 CV of wash buffer 2 (25 mM HEPES pH 7.0, 250 mM NaCl, 30 mM imidazole, 5% glycerol, 0.05% DDM, 0.005% CHS, and ligand). A_2A_AR was eluted with an elution buffer (50 mM HEPES pH 7.0, 250 mM NaCl, 300 mM imidazole, 5% glycerol, 0.05% DDM, 0.005% CHS, and ligand). The eluted protein was exchanged into a final buffer 25 mM HEPES pH 7.0, 75 mM NaCl, 0.05% DDM, 0.005% CHS, and ligand), using a PD-10 desalting column (Cytiva) for use in all further experiments. All buffers were prepared with a saturating concentration of the required ligand. For samples with LMNG/CHS, solubilization buffer was made with 0.5% LMNG and 0.025% CHS, and wash buffers, elution buffer was made with 0.05% LMNG and 0.0025% CHS, and the final buffer had 0.03% LMNG and 0.0015% CHS.

#### Expression and Purification of MSP1D1

MSP1D1 was produced for nanodisc assembly following previously described protocols.^69,70^ The MSP1D1 plasmids were transformed in BL21(DE3) cells and a single colony was used to inoculate 5 mL of LB broth supplemented with 50 μg/mL of kanamycin and grown at 37 °C overnight. The 5 mL cultures were then subsequently used to inoculate 1 L of terrific broth (TB) media and grown at 37 °C until the OD_600nm_ reached 0.6-0.8. At this point, 1 mM isopropyl β-D-1-thiogalactopyranoside (IPTG) was used to induce MSP1D1 expression, which was allowed to continue for 4 h at 30 °C. The cells were then harvested by centrifugation, and cell pellets were frozen and stored at -80 °C for later purification.

Cell pellets containing expressed MSP1D1 were thawed and resuspended in lysis buffer (50 mM Tris-HCl, pH 8.0, 500 mM NaCl, 1 mM EDTA, 1% (v/v) Triton X-100, and in-house protease inhibitor solution) and lysed by passing through a cell disruptor (Pressure Biosciences) at 20 kPSI. The cell lysate was spun down at 20,000 x g for 45 min at 4 °C and the resulting supernatant was incubated with Ni-NTA resin (pre-equilibrated with wash buffer 1 (50 mM Tris-HCl, pH 8.0, 500 mM NaCl, and 1% (v/v) Triton X-100)) for 2 h at 4°C. The Ni-NTA resin was then collected and washed with 5 CVs of wash buffer 1, 5 CVs of wash buffer 2 (50 mM Tris-HCl, pH 8.0, 500 mM NaCl, and 50 mM sodium cholate), 5 CVs of wash buffer 3 (50 mM Tris-HCl, pH 8.0, and 500 mM NaCl) and finally 5 CVs of wash buffer 4 (50 mM Tris-HCl, pH 8.0, 500 mM NaCl, and 20 mM imidazole). The MSP1D1 was eluted with the help of an elution buffer (50 mM Tris-HCl, pH 8.0, 500 mM NaCl, and 500 mM imidazole). The eluted MSP1D1 was dialyzed against a dialysis buffer (50 mM Tris-HCl, pH 8.0, 20 mM NaCl, and 0.5 mM EDTA) using a 10 kDa MWCO snake-skin dialysis tubing. Subsequently, MSP1D1 was incubated for 16 h with TEV protease at a ratio of 1:100 (TEV: MSP1D1) (w/w) at 4 °C. Following this, TEV-free MSP1D1 was isolated by incubating the mixture with Ni-NTA resin and collecting the flow-through. The MSP1D1 was dialyzed against a storage buffer (20 mM Tris-HCl, pH 8.0, 100 mM NaCl, and 0.5 mM EDTA) for 4 h at 4 °C. The purified MSP1D1 was then concentrated to a final concentration of 1 mM, aliquoted and flash frozen and stored at -80 °C for future use.

#### Nanodisc assembly

Lipid nanodiscs assembly was carried out according to previously published protocols that were optimized for multiple lipid compositions.^19^ 100 mM stock solutions of all lipids (POPC, POPS, POPE, POPG and PI(4,5)P_2_) were prepared in a cholate buffer (25 mM Tris-HCl, pH 8.0, 150 mM NaCl, and 200 mM sodium cholate). For cholesterol-containing nanodiscs, phospholipid stocks in chloroform were co-dried with cholesterol to make a lipid film and then vacuum dried for 16 h. The dried lipid film was resuspended in cholate buffer to a final phospholipid concentration of 100 mM. Nanodiscs were assembled by mixing 27 µM of purified A_2A_AR with purified MSP1D1 and detergent-solubilized lipids at a molar ratio of 1:5:250 respectively. The mixture was incubated for 2 h at 4 °C and then incubated overnight with pre-washed bio-beads at 4 °C. After the overnight incubation, the bio-beads were removed, and the resulting mixture was incubated with Ni-NTA resin for 24 h at 4 °C. Subsequently, the Ni-NTA resin was collected and washed with 2 CVs of wash buffer (50 mM HEPES, pH 7.0, 150 mM NaCl, and 10 mM imidazole). Nanodiscs containing A_2A_AR were eluted with elution buffer (50 mM HEPES, pH 7.0, 150 mM NaCl, 300 mM imidazole and ligand) and then exchanged into a final buffer (25 mM HEPES pH 7.0, 75 mM NaCl, 100 µM TFA and ligand) using a PD-10 desalting column. All ligand containing buffers were prepared with saturating amounts of ligand.

#### Membrane Fluidity Measurements

Nanodiscs containing POPC, POPE:POPS (70:30, molar ratio), POPC:POPS (70:30, molar ratio), and POPS were prepared and mixed with Laurdan (Tocris Bioscience) at a molar ratio of 500:1 (total lipids:Laurdan), and the mixture was incubated in the dark at 25 °C for 1 h. Following this incubation, excess Laurdan was removed via buffer exchange using a PD MiniTrap G-25 column (Cytiva) equilibrated with buffer (25 mM HEPES pH 7.0 and 75 mM NaCl). Fluorescence emission curves for Laurdan embedded in nanodiscs were measured in a Cary Eclipse spectrofluorometer, using an excitation wavelength of 366 nm and observing emission between 400 nm and 600 nm.

#### cAMP accumulation assays

The plasmid encoding the wild-type human A_2A_ adenosine receptor was transfected into CHO-K1 cells (ATCC product CCL-61) using lipofectamine 2000. The cell line was authenticated by the manufacturer and determined to be free from mycoplasma via a PCR based assay, agar culture method, and Hoechst DNA stain method. 24 h after transfection, cells were detached and grown in 96-well plates in medium containing equal volume of DMEM and F12 supplemented with 10% fetal bovine serum, 100 Units/ml penicillin, 100 μg/ml streptomycin, and 2 μmol/ml glutamine. After growing for 24 h, culture media was removed, and cells were washed twice with PBS. Cells were then treated with assay buffer containing rolipram (10 μM) and adenosine deaminase (3 units/ml) for 30 min followed by addition of agonist and incubated for 20 min. The reaction was terminated upon removal of the supernatant, and addition of 100 μL Tween-20 (0.3%). Intracellular cAMP levels were measured with an AlphaScreen cAMP assay kit (PerkinElmer, Catalog Number: 6760635D) following the manufacture’s protocol.

#### NMR experiments

Nanodisc samples containing A_2A_AR were concentrated to ∼200 μM in 280 μL using a Vivaspin-6 concentrator with a 30 kDa MWCO, and 20 μL of D2O was mixed in gently into the sample. ^19^F-NMR experiments were recorded on a Bruker Avance III HD spectrometer operating at 600 MHz ^1^H nutation frequency using Topspin 3.6.2 and equipped with a Bruker 5-mm BBFO probe. ^19^F NMR data was recorded at three different temperatures, 280 K, 298 K and 310 K. Temperatures were calibrated from a standard sample of 4% methanol in D_4_-MeOH.

1-dimensional ^19^F NMR data were recorded with a data size of 32k complex points, with an acquisition period of 360 ms, a 120 μs dwell time and 0.3 s recycle delay for a total experimental time of ∼ 3.5 h per experiment. All ^31^P NMR experiments were carried out with the recycle delay of 0.3 s, 2k scans, and an acquisition time of 900 ms for a total experiment time of 42 min per experiment.

### QUANTIFICATION AND STATISTICAL ANALYSIS

#### NMR data analysis

All NMR data were processed and analyzed in Topspin 4.0.8 (Bruker Biospin). All 1-dimensional ^19^F-NMR data were processed identically. The data were zero-filled to 64k points and multiplied by an exponential window function with 40 Hz line broadening prior to Fourier transformation. ^19^F spectra were referenced to the signal from trifluoroacetic acid (TFA) at -75.8 ppm, which was set to 0 ppm. Deconvolution of the ^19^F-NMR data followed previously published procedures and was done with MestreNova version 14.1.1-24571 (Mestrelab Research). For each spectrum, the between the experiment data and sum of the deconvoluted signals was assessed to check the quality of deconvolution. The relative population of the different A_2A_AR conformational states were calculated as a ratio of integrated area of the each deconvoluted peak to the total integral of all the signals from 6 ppm to 15.5 ppm.

For ^31^P NMR experiments prior to Fourier transformation, ^31^P data were zero-filled to 64k points and multiplied by an exponential window function with 50 Hz line broadening. All ^31^P NMR data were analyzed identically in Topspin 4.0.8 (Bruker Biospin).

#### Membrane Fluidity Data Analysis

The obtained emission spectra were normalized to the corresponding emission maxima. In accordance with earlier protocols from Yu et al.,^71^ the generalized polarization (GP) of Laurdan in the nanodiscs was calculated using the formula:

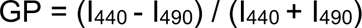

where I_440_ and I_490_ are the fluorescence emission intensities at 440 nm and 490 nm, respectively. All GP values reported are the mean ± s.e.m. (standard error of mean) for 3 independent measurements.

#### Analysis of cAMP accumulation data

Functional parameters for cAMP accumulation assay were calculated using Prism 9.10 software (GraphPad Software, San Diego, CA). Data from functional assays were normalized by considering the signal levels of cells alone to be 0%, and stimulation by the full agonist NECA was considered to be 100%. All reported values are mean ± s.e.m. for three independent experiments.

---

**Figure S1:**
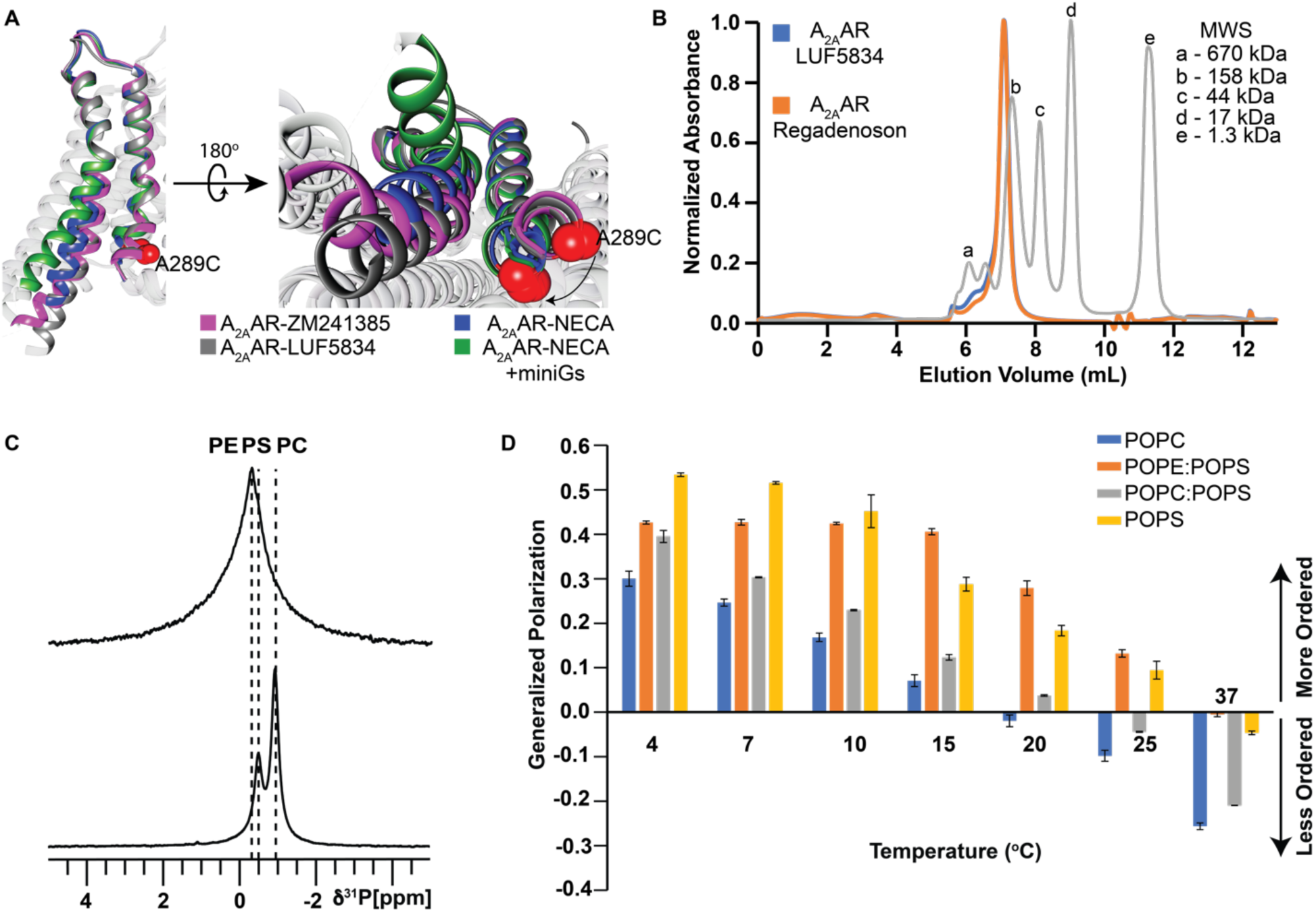
Biophysical characterization and membrane fluidity measurements of nanodiscs of varied lipid compositions. Related to Figures 1, 2 and 3. (A) Structures of A_2A_AR in complex with antagonist ZM241385 (pink ribbon, PDB ID: 3EML), partial agonist LUF5834 (grey ribbon, PDB ID: 7ARO), agonist NECA (blue ribbon, PDB ID: 2YDV), and agonist NECA + mini-Gs (green ribbon, PDB ID: 5G53). For clarity, only helices 6 and 7 are shown here. (Right) Intracellular view of helices 6 and 7, highlighting structural changes on helix 6 and 7. Position 289 is used as a site for ^19^F-NMR labelling and shown as a red sphere. (B) Analytical size exclusion chromatograms of A_2A_AR in complex with Regadenoson (orange) and LUF5834 (blue) in POPC:POPS (70:30) nanodiscs. The grey chromatogram represents molecular weight standards with the molecular weights indicated on the corresponding peaks. (C) 1-dimensional ^31^P-NMR of nanodiscs containing A_2A_AR[A289C^TET^] and POPE:POPS (70:30, molar ratio) or A_2A_AR[A289C^TET^] and POPC:POPS (70:30, molar ratio). Signals assigned to POPE, POPS, and POPC are indicated by the vertical dashed lines with chemical shifts consistent with the manufacturer’s reported values. The relative intensities of the two observed signals were qualitatively consistent with the expected lipid ratios in the nanodisc samples. (D) Laurdan-based fluidity measurements of membrane lipids in nanodiscs containing POPC, POPE:POPS (70:30), POPC:POPS(70:30), and POPS across a range of temperature from 4 °C to 37 °C. Generalized polarization values are plotted as the mean ± s.e.m. for three independent trials.

**Figure S2.**
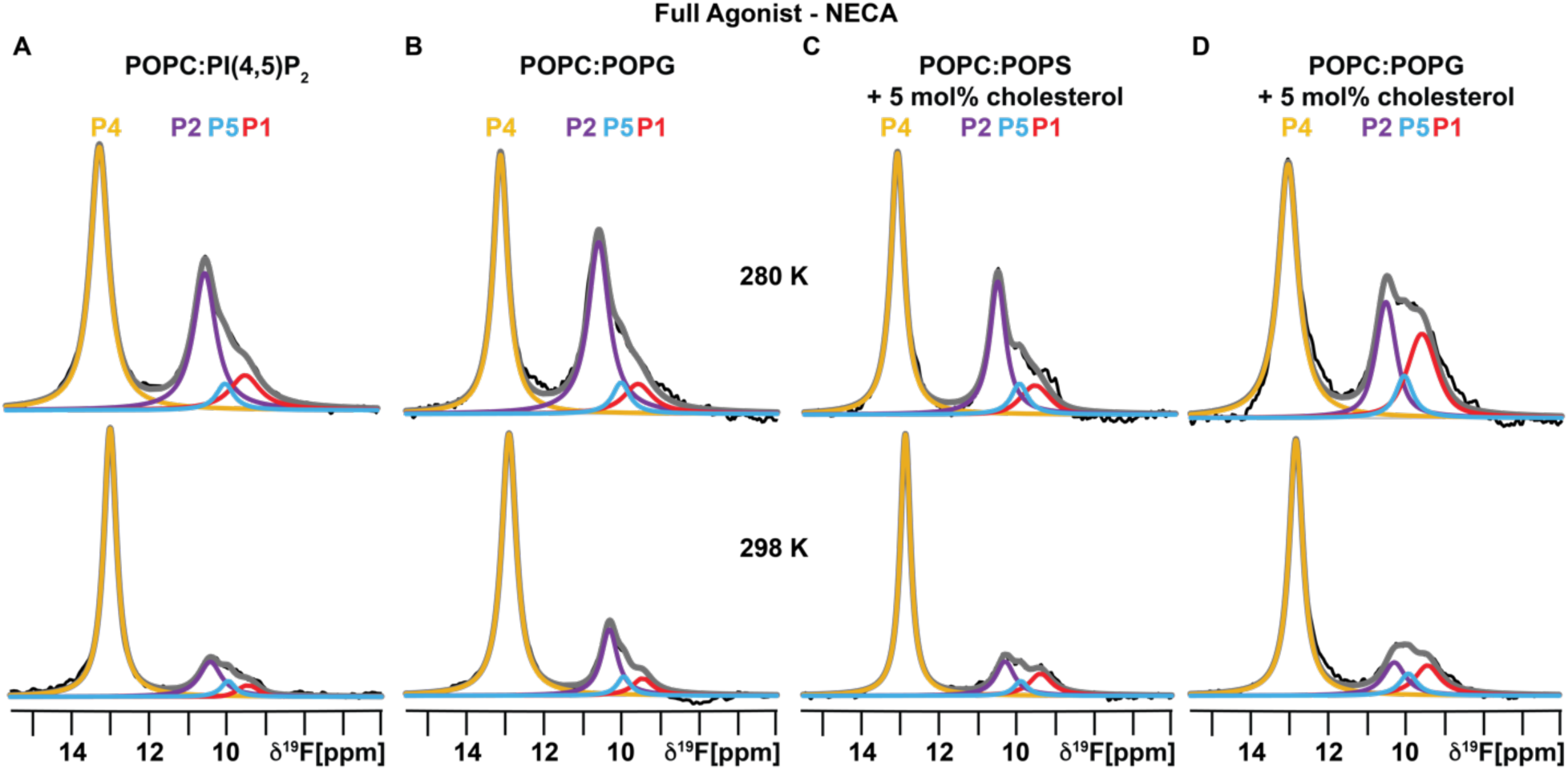
Variable temperature ^19^F-NMR spectra of agonist-bound A_2A_AR reconstituted in nanodiscs of varied lipid composition as a response to temperature. Related to Figure 1. (A – D) 1-dimensional ^19^F NMR spectra of A_2A_AR-NECA reconstituted in (A) POPC:PI(4,5)P_2_ (95:5, molar ratio), (B) POPC:POPG (70:30, molar ratio), (C) POPC:POPS:Cholesterol (70:30:5 mol%), and (D) POPC:POPG:Cholesterol (70:30:5 mol%) recorded at 280 K and 298 K.

**Figure S3:**
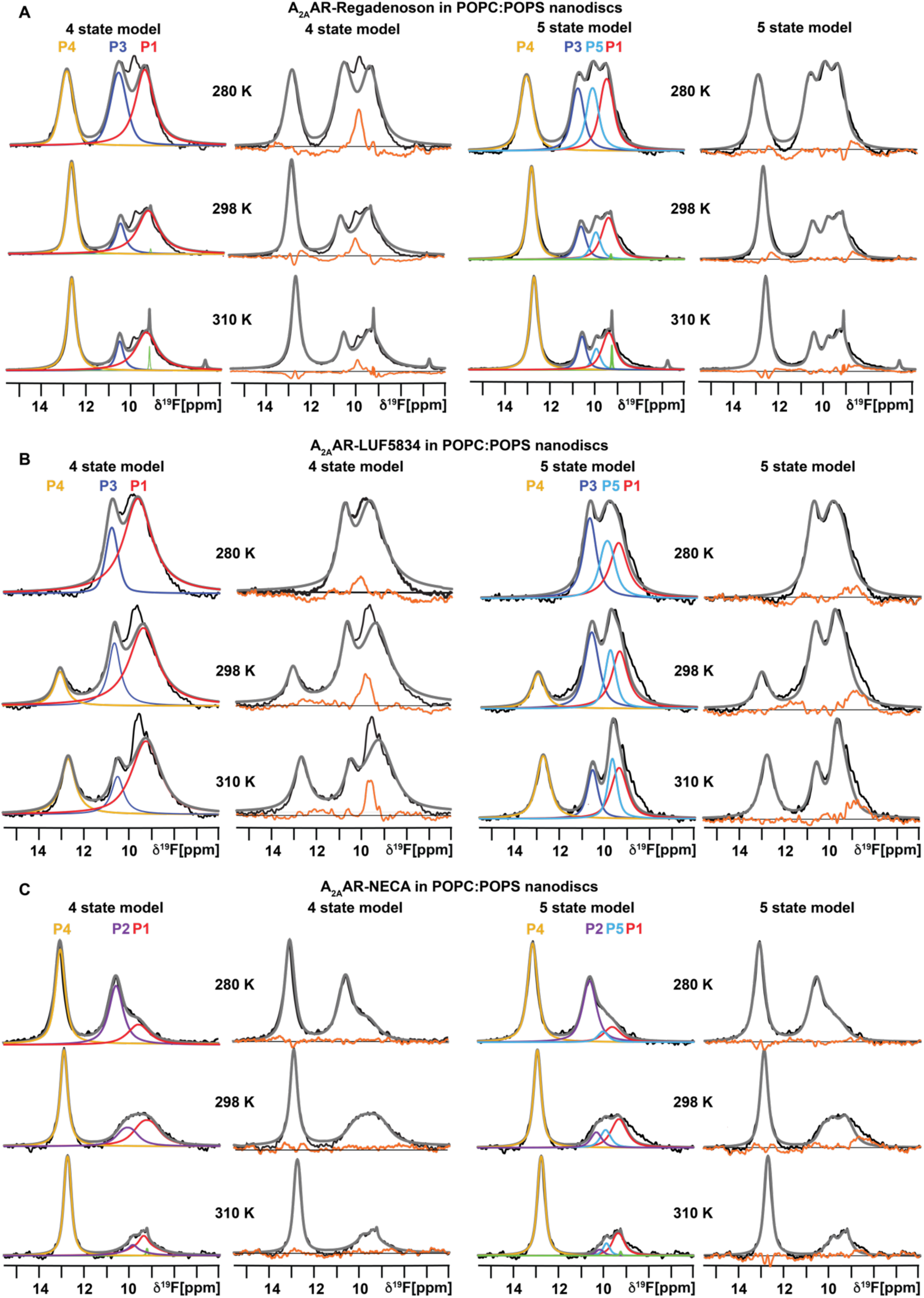
^19^F-NMR data fitting to different models. Related to Figure 1 and 2. (A – C) ^19^F NMR data of (A) A_2A_AR-Regadenoson, (B) A_2A_AR-LUF5834, and (C) A_2A_AR-NECA in nanodiscs containing POPC:POPS (70:30, molar ratio) recorded at 280 K, 298 K, and 310 K. In each panel, the data on the left half are fit to a 4-state model and the date on the right half are fit to a 5-state model, as indicated. Other presentation details are same as Figure 1.

**Figure S4:**
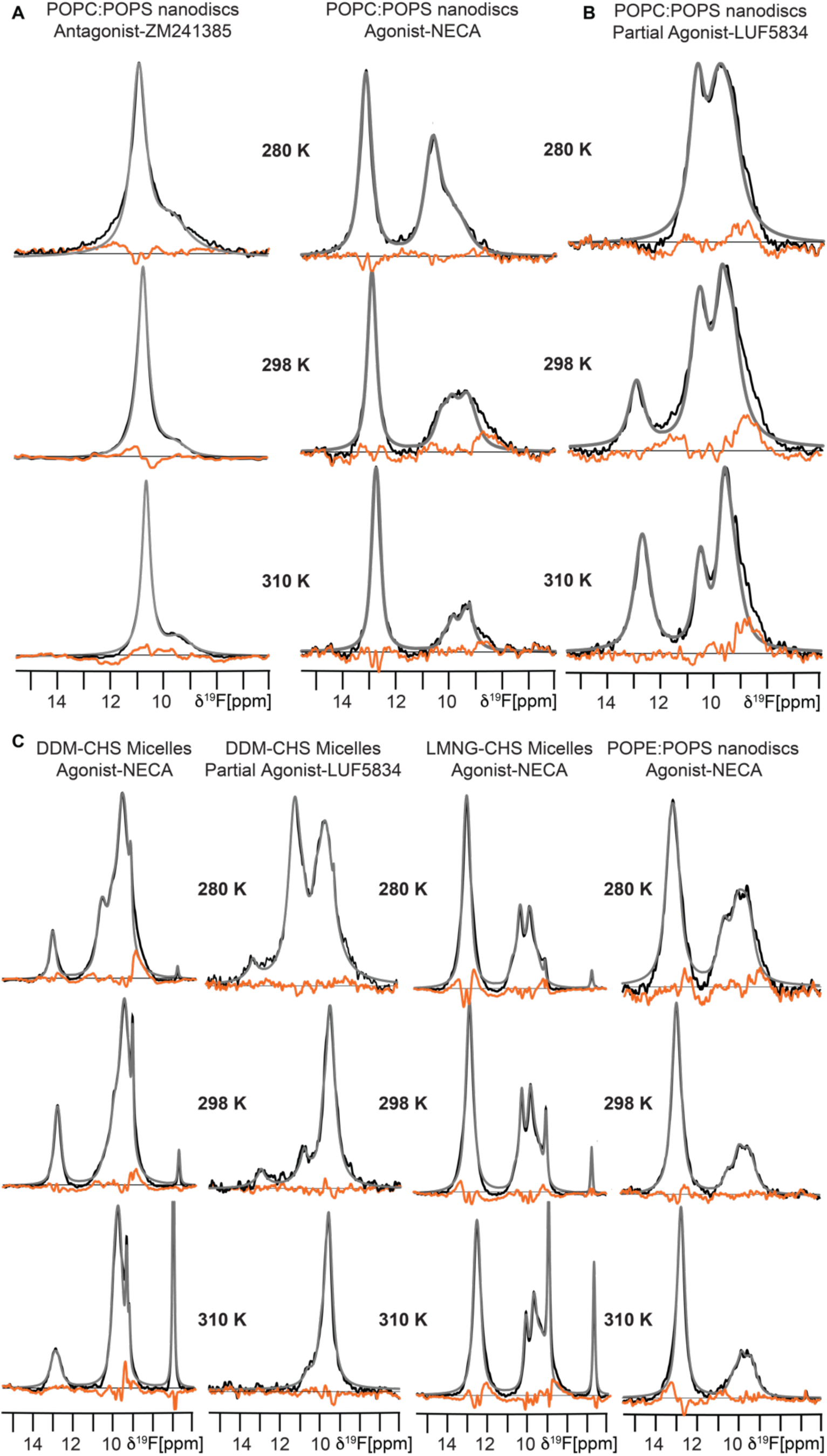
Residual differences between the raw data and summation of the individually fit components from Figures 1, 2, and 3. (A – C) The experimental data are shown in black, the grey lines superimposed on the spectra are the total sums of the individual deconvolutions, and the orange line is the calculated difference between the grey and black lines.

**Table S1:**
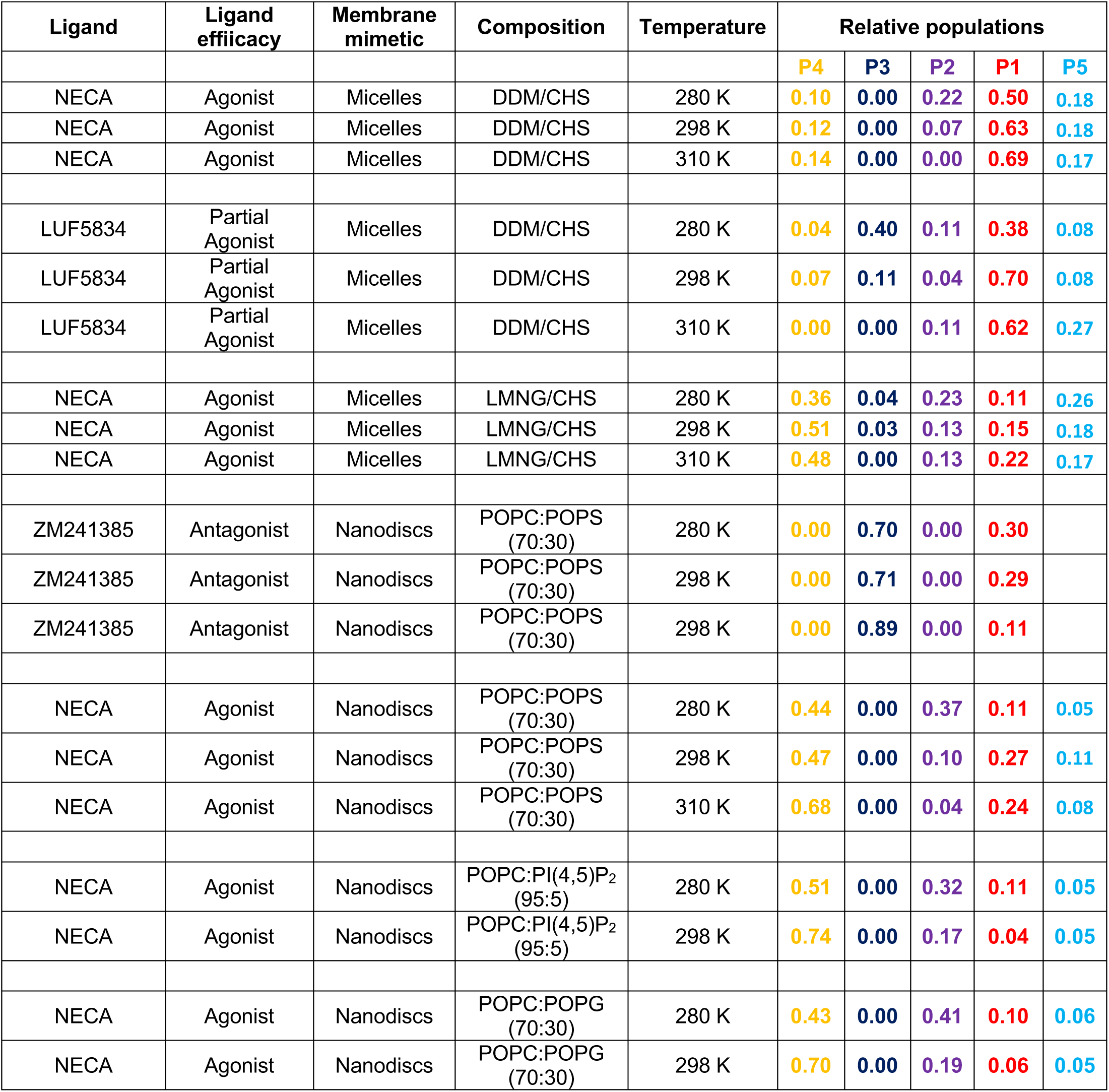
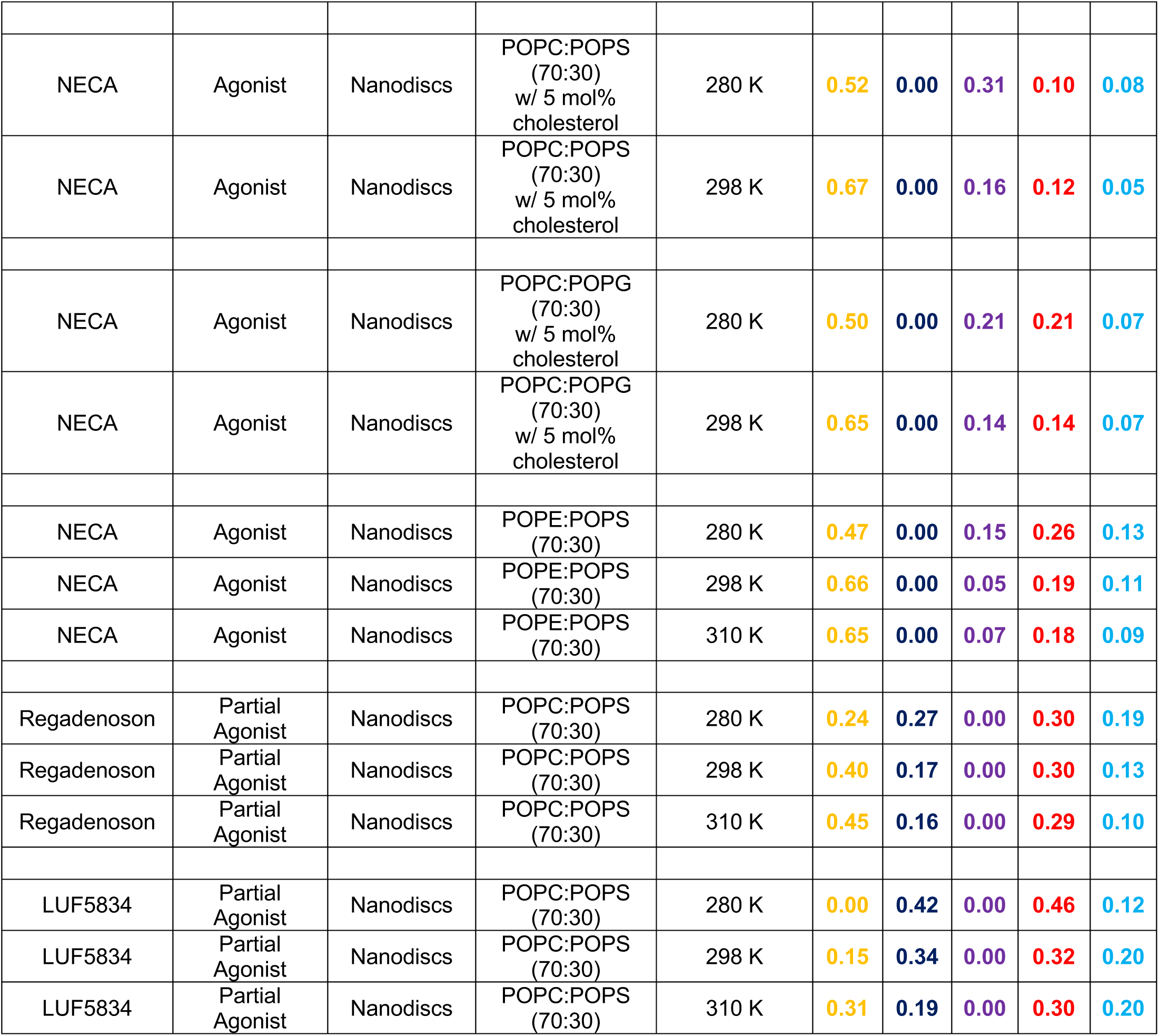
Relative populations of the A_2A_AR conformational states observed in ^19^F-NMR. Related to Figure 1, 2, and 3. The relative populations of the A_2A_AR conformational states as observed in the ^19^F-NMR spectra are tabulated below. The value reported for each conformation is a ratio of the integrated area of the specific deconvoluted peak to the total integral of all signals from 6 ppm to 15.5 ppm, excluding any signals from free TET.

| Ligand | Ligand efficacy | Membrane mimetic | Composition | Temperature | Relative populations |  |  |  |  |
| --- | --- | --- | --- | --- | --- | --- | --- | --- | --- |
|  |  |  |  |  | P4 | P3 | P2 | P1 | P5 |
| NECA | Agonist | Micelles | DDM/CHS | 280 K | 0.10 | 0.00 | 0.22 | 0.50 | 0.18 |
| NECA | Agonist | Micelles | DDM/CHS | 298 K | 0.12 | 0.00 | 0.07 | 0.63 | 0.18 |
| NECA | Agonist | Micelles | DDM/CHS | 310 K | 0.14 | 0.00 | 0.00 | 0.69 | 0.17 |
| LUF5834 | Partial Agonist | Micelles | DDM/CHS | 280 K | 0.04 | 0.40 | 0.11 | 0.38 | 0.08 |
| LUF5834 | Partial Agonist | Micelles | DDM/CHS | 298 K | 0.07 | 0.11 | 0.04 | 0.70 | 0.08 |
| LUF5834 | Partial Agonist | Micelles | DDM/CHS | 310 K | 0.00 | 0.00 | 0.11 | 0.62 | 0.27 |
| NECA | Agonist | Micelles | LMNG/CHS | 280 K | 0.36 | 0.04 | 0.23 | 0.11 | 0.26 |
| NECA | Agonist | Micelles | LMNG/CHS | 298 K | 0.51 | 0.03 | 0.13 | 0.15 | 0.18 |
| NECA | Agonist | Micelles | LMNG/CHS | 310 K | 0.48 | 0.00 | 0.13 | 0.22 | 0.17 |
| ZM241385 | Antagonist | Nanodiscs | POPC:POPS (70:30) | 280 K | 0.00 | 0.70 | 0.00 | 0.30 |  |
| ZM241385 | Antagonist | Nanodiscs | POPC:POPS (70:30) | 298 K | 0.00 | 0.71 | 0.00 | 0.29 |  |
| ZM241385 | Antagonist | Nanodiscs | POPC:POPS (70:30) | 298 K | 0.00 | 0.89 | 0.00 | 0.11 |  |
| NECA | Agonist | Nanodiscs | POPC:POPS (70:30) | 280 K | 0.44 | 0.00 | 0.37 | 0.11 | 0.05 |
| NECA | Agonist | Nanodiscs | POPC:POPS (70:30) | 298 K | 0.47 | 0.00 | 0.10 | 0.27 | 0.11 |
| NECA | Agonist | Nanodiscs | POPC:POPS (70:30) | 310 K | 0.68 | 0.00 | 0.04 | 0.24 | 0.08 |
| NECA | Agonist | Nanodiscs | POPC:PI(4,5)P<sub>2</sub> (95:5) | 280 K | 0.51 | 0.00 | 0.32 | 0.11 | 0.05 |
| NECA | Agonist | Nanodiscs | POPC:PI(4,5)P<sub>2</sub> (95:5) | 298 K | 0.74 | 0.00 | 0.17 | 0.04 | 0.05 |
| NECA | Agonist | Nanodiscs | POPC:POPG (70:30) | 280 K | 0.43 | 0.00 | 0.41 | 0.10 | 0.06 |
| NECA | Agonist | Nanodiscs | POPC:POPG (70:30) | 298 K | 0.70 | 0.00 | 0.19 | 0.06 | 0.05 |
| NECA | Agonist | Nanodiscs | POPC:POPS<br>(70:30)<br>w/ 5 mol%<br>cholesterol | 280 K | 0.52 | 0.00 | 0.31 | 0.10 | 0.08 |
| NECA | Agonist | Nanodiscs | POPC:POPS<br>(70:30)<br>w/ 5 mol%<br>cholesterol | 298 K | 0.67 | 0.00 | 0.16 | 0.12 | 0.05 |
| NECA | Agonist | Nanodiscs | POPC:POPG<br>(70:30)<br>w/ 5 mol%<br>cholesterol | 280 K | 0.50 | 0.00 | 0.21 | 0.21 | 0.07 |
| NECA | Agonist | Nanodiscs | POPC:POPG<br>(70:30)<br>w/ 5 mol%<br>cholesterol | 298 K | 0.65 | 0.00 | 0.14 | 0.14 | 0.07 |
| NECA | Agonist | Nanodiscs | POPE:POPS<br>(70:30) | 280 K | 0.47 | 0.00 | 0.15 | 0.26 | 0.13 |
| NECA | Agonist | Nanodiscs | POPE:POPS<br>(70:30) | 298 K | 0.66 | 0.00 | 0.05 | 0.19 | 0.11 |
| NECA | Agonist | Nanodiscs | POPE:POPS<br>(70:30) | 310 K | 0.65 | 0.00 | 0.07 | 0.18 | 0.09 |
| Regadenoson | Partial<br>Agonist | Nanodiscs | POPC:POPS<br>(70:30) | 280 K | 0.24 | 0.27 | 0.00 | 0.30 | 0.19 |
| Regadenoson | Partial<br>Agonist | Nanodiscs | POPC:POPS<br>(70:30) | 298 K | 0.40 | 0.17 | 0.00 | 0.30 | 0.13 |
| Regadenoson | Partial<br>Agonist | Nanodiscs | POPC:POPS<br>(70:30) | 310 K | 0.45 | 0.16 | 0.00 | 0.29 | 0.10 |
| LUF5834 | Partial<br>Agonist | Nanodiscs | POPC:POPS<br>(70:30) | 280 K | 0.00 | 0.42 | 0.00 | 0.46 | 0.12 |
| LUF5834 | Partial<br>Agonist | Nanodiscs | POPC:POPS<br>(70:30) | 298 K | 0.15 | 0.34 | 0.00 | 0.32 | 0.20 |
| LUF5834 | Partial<br>Agonist | Nanodiscs | POPC:POPS<br>(70:30) | 310 K | 0.31 | 0.19 | 0.00 | 0.30 | 0.20 |

